# Importance of Leaf Age in Grapevines Under Salt Stress

**DOI:** 10.1101/2023.04.02.535264

**Authors:** Yaniv Lupo, Kakarla Prashanth, Naftali Lazarovitch, Aaron Fait, Shimon Rachmilevitch

**Author notes:** **Correspondence:**, French Associates Institute for Agriculture and Biotechnology of Dryland, The Jacob Blaustein Institutes for Desert Research, Ben-Gurion University of the Negev, Sede-Boker Campus, 8499000, Israel.

## Abstract

The combined effects of salt stress and leaf age on photosynthesis and stomatal conductance are not fully understood. Salt stress develops at different rates in various plant organs and tissues, leading to osmotic and ionic stresses. We studied the development of salt stress (30 mM NaCl) in two grapevine rootstocks by examining the gas exchange, photochemistry, and salt accumulation of leaves at different ages. Our results indicate that leaf age strongly affected photosynthesis but had a minor effect on stomatal conductance in both control and salinity conditions. Surprisingly, salt stress lowered stomatal conductance but had little effect on photosynthesis, opposite to the leaf age effects. Reduction in photosystem II efficiency explained the age-associated decline in photosynthesis, suggesting that the light-dependent phase of photosynthesis is the first process affected by leaf aging. We also found that the sodium exclusion capacity decreased as leaves matured and was age and variety dependent, indicating the importance of leaf age when sampling for salt tolerance studies. Therefore, we suggest that sampling only young leaves is missing critical information in the evaluation of salt-tolerant traits in grapevines.

## 1. Introduction

Photosynthesis is a fundamental process for plants and one of the most researched processes in plant science (Stirbet et al., 2020). The relationship between photosynthesis and leaf age has been investigated for almost a century. Young, fully expanded leaves were found to have the highest photosynthetic rate, mature leaves with a lower rate, and old leaves with the lowest photosynthetic rate (Singh and Lal, 1935). In woody perennials, such as grapevines (*Vitis vinifera*), the photosynthesis rate is at its maximum when the leaf reaches its full size (Kriedemann et al., 1970), followed by a steady decline as the leaf matures (Intrieri et al., 1992). Similar to what was also found in deciduous trees’ photosynthetic capacity (Wilson et al., 2001). Grapevines’ maximum photosynthesis rates were found in leaves 20 to 40 days old (Poni and Intrieri, 2001; Schubert et al., 1996). The reduction of photosynthesis in aging leaves can arise from different factors such as reduction in mesophyll conductance, reduction in rubisco efficiency, or accumulation of ROS (Farquhar and Sharkey, 1982; Flexas et al., 2012; Stitt and Schulze, 1994). The relationship between leaf age and transpiration was less investigated and reached contradicting results. Regardless of leaf maturation and senescence, reduced g_s_ in old leaves can result from their basal position in the plant, exposing them to lower irradiation (Turner and Begg, 1973). However, this was not the only factor, as cotton and potato’s old leaves had lower g_s_ than the young leaves in response to drought, which was irrelevant to irradiation (Jordan et al., 1975; Vos and Oyarzún, 1987).

Salinity was estimated to affect 20%-24% of the world’s irrigated lands, and 1-1.5 Mha becomes salinized yearly (Squires and Glenn, 2011). This fraction will most likely increase in the future by increasing use of dryland for agriculture due to increasing population pressure (Metternicht and Zinck, 2003). Most crop plants are somewhat sensitive to salinity (Munns and Tester, 2008). Salt stress initially affects the plant via osmotic stress, similar to drought stress. In addition to the osmotic stress, salt affects plants via ionic and oxidative stresses (Rasool et al. 2013). The ionic phase develops slower and starts when salt-derived ions (most commonly Na^+^ and Cl^-^) reach toxic levels in plant cells. These phases usually co-occur, and it is often hard to evaluate if the response is adaptive or results from damage to the plant (Zhou-Tsang et al., 2021). Salt stress reduces grapevines’ growth rate and yield and, in severe cases, leads to plant mortality (Shani and Ben-Gal, 2005). Plants’ salt stress tolerance mechanisms include morphological and osmotic adjustments as well as salt exclusion mechanisms (whole plant, tissue, or cell levels). However, they come with energy costs to the plant (Munns and Gilliham, 2015).

Grapevines are good sodium (Na^+^) excluders, delaying Na^+^ from reaching toxic levels in their roots and leaves (Li et al., 2017; Lupo et al., 2022). Thus, chloride (Cl^-^) is more toxic for them. Cl^-^ is a beneficial macronutrient. Cl^-^ was found to increase water use efficiency (WUE) when supplied in low concentrations (Franco-Navarro et al., 2019). However, under saline conditions, Cl^-^ accumulates to toxic levels in the plant. Growing desired grape varieties on salt-tolerant rootstocks is the primary method to cope with salinity. Rootstock variety highly contributed (19%) to the variability of grapevines in response to the environment (Lavoie-Lamoureux et al., 2017). Growing the same variety on different rootstocks was found to reduce or improve its yield under long-term salt stress, depending on the rootstock (Walker et al., 2002). The selection of a rootstocks-scion combination is crucial as it also affects the scion growth, ion allocations, and yield (Gautier et al., 2020; Walker et al., 2004).

Leaf age and its relationship to photosynthesis have been investigated for almost a century. However, the mechanisms underlying the age-associated decline in photosynthesis are not yet fully understood. To address this gap in knowledge, we examined how leaf age affects gas exchange and salt tolerance traits, such as salt exclusion, in grapevines exposed to saline conditions. We also evaluated the responses of two moderate salt-tolerant rootstocks (Bettiga, 2003) from different genetic backgrounds to salt stress. We hypothesized that salt concentration will increase with leaf age and that it will be correlated with decreased leaf function (e.g., CO_2_ fixation rate). This study aims to improve our understanding of how salt stress develops in a maturing leaf, how leaf age affects its function, and add to the knowledge on salt tolerance of different grapevine rootstocks.

## 2. Materials and Methods

### 2.1 Experimental design

The experiment was conducted in a net house at the Sde Boker campus of the Ben-Gurion University of the Negev (31° N, 35° E). Two commercially used grapevine rootstocks were used: Paulsen 1103 (V. berlandieri * V. rupestris; 1103P) and Sélection Oppenheim 4 (V. berlandieri * V. riparia; SO4). Four weeks old cuttings were planted on May 3, 2020, in 20-liter pots filled with sand (∼85% sand, ∼10% silt, and ∼5% clay) and covered with a plastic lid. The pots were set in 3 blocks facing east-west, and nine replicates were used for each rootstock treatment combination. Plants were continuously pruned to allow one main shoot to grow, and flowers were removed to minimize variables. The vines were trellised to maximize leaf light exposure (see S fig. 1). The plants were irrigated using a drip irrigation system (1.2 l/h pressure compensating drippers) connected directly to the final fertilizer and salt solutions tanks. The plants were excessively irrigated every 2 days, maintaining a leaching fraction of 33% to prevent water deficiency or NaCl accumulation in the root system. A 16.7% Hoagland solution was used for fertilization to supply 40 ppm of nitrogen (Hoagland and Arnon, 1950). Two salinity levels were used: 1. Freshwater + fertilizer (0.7 dS m^-1^; control); 2. Freshwater + fertilizer + added salt (30 mM NaCl, 4 dS m^-1^). Salt treatment began on June 3, 2020. A pipe filled with rock wool was connected to the bottom of each pot to improve sand aeration and leaching (Ben Gal and Shani, 2002). All plants were harvested at the end of the experiment, August 10, 2020, and measured for shoot mass and stem diameter. Shoots (stems + leaves) were cut 5 cm from the soil surface, oven-dried at 65 °C for 72 hours, and then measured for dry mass. The stem diameter was measured 20 cm from the soil surface with a digital caliper.

### 2.2 Gas exchange measurements

Gas-exchange measurements were taken with the LI-6800 portable photosynthesis system (LI-COR, Lincoln, NE). Midday measurements were done every week between 12 pm to 2 pm on the same leaves. Chamber parameters were set to 400 ppm CO2, 1000 μmol m^-2^ s^-1^ PFD, 35 °C air temperature and 50% RH. Diurnal measurements were done on young (21 days) and mature (63 days) leaves at 6 am, 8 am, 10 am, 12 pm, 2 pm, 4 pm, and 6 pm. Chamber parameters on the diurnal trend measurements were set to be similar to the hourly environmental conditions. Diurnal measurements were taken 42 and 43 days after treatment started for SO4 and 1103P, respectively.

### 2.3 Soluble ions, elemental analysis, and Osmolality measurements

The youngest unfolded leaves were marked for ion analysis every two weeks. At the end of the experiment, leaf blades were collected from marked leaves, washed with distilled water, and oven dried for 72 hours at 65 °C. Leaf tissue was grounded to powder using RETCH-mill, then 50 mg powder was collected in 15 ml plastic tubes for ion analysis. Cl^-^ was extracted in distilled water (5 ml) and analyzed by a silver ion titration Cl^-^ analyzer (M926, Sherwood Scientific Ltd., Cambridge, UK). Na^+^ was extracted in 0.5 M nitric acid (5 ml) and analyzed by an inductively coupled plasma–optical emission spectrometer (ICP OES; Varian 720 ES, Varian Inc, CA, US) following (Munns et al., 2010). Samples of 3.5 g were analyzed for carbon and nitrogen by an organic elemental analyzer (Flash 2000, Thermo Scientific, MA, US). Young (21 days) and mature (63 days) leaves were sampled for osmolality on August 5 at two-time points: morning (6 am) and midday (12 pm). The leaves were immediately placed in 2 ml tubes and frozen with liquid nitrogen. Leaf sap was extracted using a centrifuge at 18,000 rpm for 2 min. Then 10 μl of leaf sap was used for osmolality measurement using a vapor pressure osmometer (Vapro® 5520, Wescor, US).

### 2.4 Statistical analysis

One-way ANOVA was performed to determine statistically significant differences between leaf ages within each variety-treatment combination. Two-way ANOVA was performed to determine statistically significant differences between leaf age-treatment combinations within each time point/variety or to determine statistically significant differences between variety-treatment combinations within each leaf age. Tukey HSD was used as a post hoc test. Student’s t-test was performed to determine statistically significant differences between treatments within each variety-leaf age combination or between varieties for specific comparisons. Data visualization and statistical analysis were performed using R (4.2.1).

## 3. Results

### 3.1 Gas exchange of leaves at different ages

1103P’s stomatal conductance (g_s_) peaked when leaves were 28 days old under the control treatment. Still, there were no significant differences between the leaf ages except from 63 days old leaves (Fig. 1a). Under the salt treatment, 1103P’s g_s_ was significantly higher when the leaves were 21 days old compared only to 14 days old leaves. There were no significant differences between different aged leaves from day 21 to the end of the experiment. SO4’s g_s_ peak, under the control treatment, was a week earlier than 1103P when its leaves were 21 days old (Fig. 1b). SO4’s g_s_ significantly reduced when its leaves were 56 days old and older. Under the salt treatment, SO4’s g_s_ peak was observed in 56 days old leaves, which was significantly higher compared to 14 and 28 days old leaves. There were no significant differences in g_s_ between 1103P and SO4 under the control treatment in all leaf ages (data not shown). 1103P had lower g_s_ under the salt treatment compared to the control when its leaves were 14, 28, 35, 42, and 70 days old. SO4 had lower g_s_ under the salt treatment compared to the control when its leaves were 14, 21, 28, 35, 42, and 49 days old.

**Figure 1.**
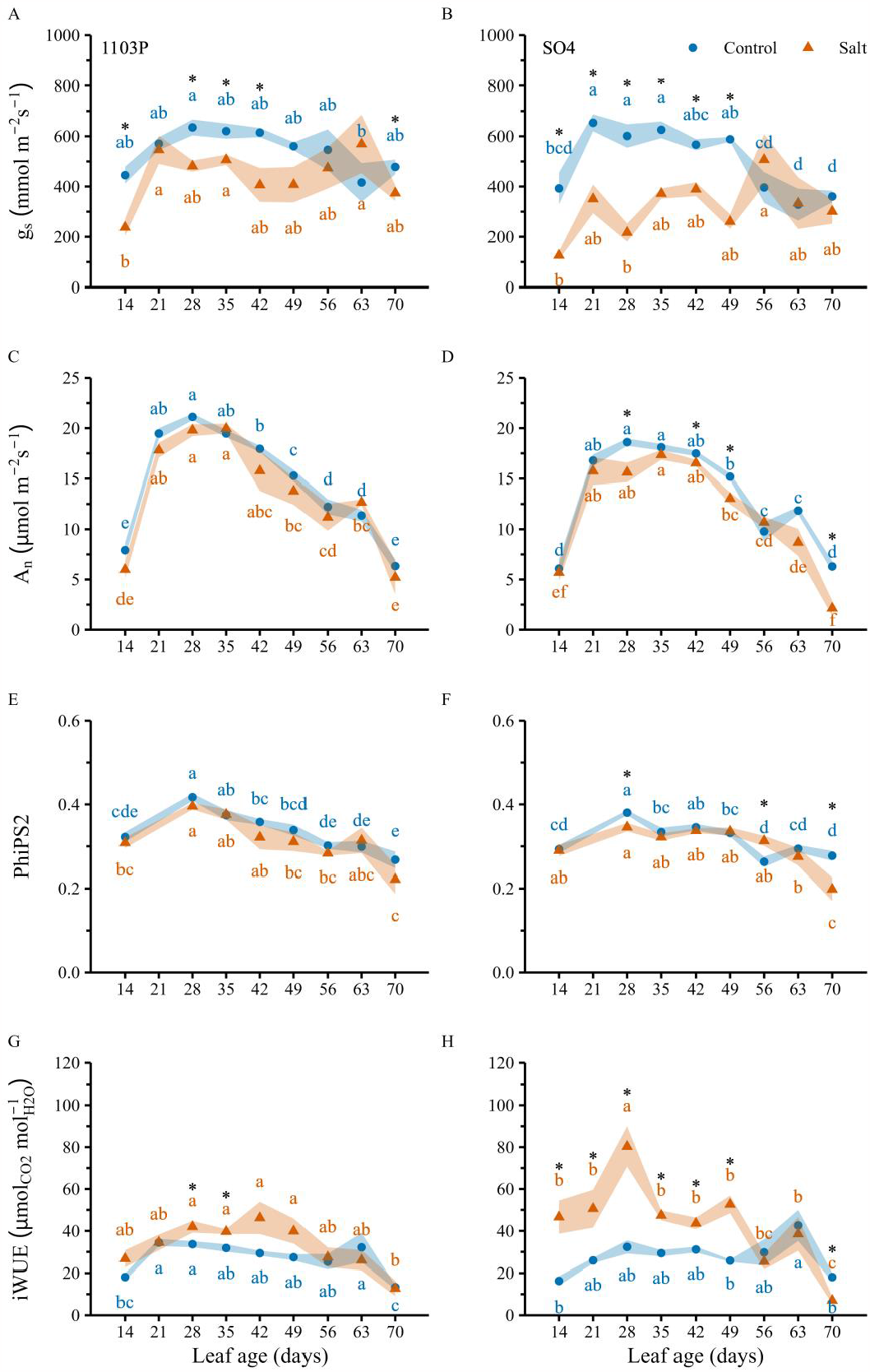
Mean gas exchange and PSII efficiency in different leaf ages of 1103P (left) and SO4 (right) under control (blue circles) and salt (orange triangles) treatments. (a) 1103P’s stomatal conductance (g_s_). (b) SO4’s g_s_. (c) 1103P’s CO2 assimilation rate (A_n_). (d) SO4’s A_n_. (e) 1103P’s Photosystem II efficiency (PhiPS2). (f) SO4’s PhiPS2. (g) 1103P’s intrinsic water use efficiency (A_n_/g_s_; iWUE). (h) SO4’s iWUE. Background color represents standard error. Different letters represent significant differences between leaf ages within each treatment, one-way ANOVA, P < 0.05. Asterisks represent significant differences between treatments within each leaf age, student’s t-test, P < 0.05. n = 6.

The CO2 assimilation rate (A_n_) of both 1103P and SO4 increased as the leaves matured and reached a peak when leaves were 28 days old under the control treatment (Fig. 1c, d). Significant reductions in A_n_ from the peak were observed when the leaves were 42 and 49 days old for 1103P and SO4, respectively. The reductions in A_n_ continued to the end of the experiment when leaves were 70 days old, with an expiation at 63 days. 1103P had higher A_n_ compared to SO4 under the control treatment when leaves were 28 days old (student’s t-test, P < 0.05; data not shown). At that age, SO4 also had lower A_n_ under the salt treatment compared to the control. SO4 had higher A_n_ under the salt treatment, compared to the control, when its leaves were 42 and 49 days old. A peak in photosystem II (PSII) efficiency (PhiPS2) was observed when leaves were 28 days old (Fig. 1e, f), similar to the peak in A_n_. The PhiPS2 was higher in 28 days old leaves compared to 14 days old leaves under the control treatment for 1103P and SO4 and under the salt treatment only for 1103P’s leaves. Significant reductions in PhiPS2, compared to the peak, were observed from 42 and 56 days old leaves in 1103P and SO4, respectively, under the control treatment. Under the salt treatment, reductions in PhiPS2 were observed when leaves were 56 and 63 days or older in 1103P and SO4, respectively.

Leaf age had little to no effect on the intrinsic water use efficiency (iWUE, A_n_/g_s_) under the control treatment for 1103P and SO4 (Fig. 1g, h). Leaf age significantly affected the iWUE only when the leaves were 14 and 70 days old or 63 days old for 1103P and SO4, respectively. In several leaf ages, the iWUE was increased under the salt treatment (more for SO4 than 1103P). 1103P had higher iWUE under the salt treatment compared to control when its leaves were 28 and 35 days old (Fig. 1g). SO4 had higher iWUE under the salt treatment compared to control when its leaves were 14, 21, 28, 35, 42, 49 and 70 days old (Fig. 1h).

### 3.2 Diurnal gas exchange of young and mature leaves

1103P’s and SO4’s young leaves started the day with higher g_s_ under the control treatment compared to the salt treatment and the mature leaves (Fig. 2a, b, table 1). SO4’s g_s_ peak was at 8 am, while 1103P’s peak was at 12 pm. 1103P’s young leaves maintained high g_s_ during the day, significantly lowering their g_s_ at 2 pm and 6 pm under the salt and control treatments, respectively (Fig. 2a, table 1). 1103P’s mature leaves reduced their g_s_ earlier than the young leaves under the control treatment, with a significant reduction at 2 pm. SO4’s young leaves reduced their g_s_ earlier compared to 1103P, with significant reductions at 10 am and 12 pm under the salt and control treatments, respectively (Fig. 2b, table 1). SO4’s mature leaves also reduced their g_s_ early, with significant reductions at 12 pm under both treatments. 1103P young leaves had higher g_s_ compared to SO4 young leaves at midday under both treatments (student’s t-test, P < 0.05; data not shown). There were no significant differences between 1103P’s and SO4’s mature leaves at midday.

**Table 1.** Two-way ANOVA analysis for the diurnal measurements of g_s_. Results are presented in figure 2a, b. Different uppercase letters represent significant differences between leaf ages and treatments within each time point. Different lowercase letters represent significant differences between time points within each leaf age-treatment combination. Tukey HSD, P < 0.05. n = 6.

| Variety | Treatment/Hour | 06:00 | 08:00 | 10:00 | 12:00 | 14:00 | 16:00 | 18:00 |
| --- | --- | --- | --- | --- | --- | --- | --- | --- |
| 1103P | Young Control | A b | A a | A ab | A a | A ab | A ab | AB c |
|  | Young Salt | B cd | AB a | A ab | AB ab | B bc | BC cd | B d |
|  | Mature Control | BC c | AB a | AB ab | A ab | B bc | B bc | A c |
|  | Mature Salt | C b | B a | B ab | B ab | B b | C b | AB b |
| SO4 | Young Control | A bc | A a | A ab | A cd | NS d | A d | NS d |
|  | Young Salt | B bc | BC a | B b | AB cd | NS cd | AB bcd | NS d |
|  | Mature Control | B cd | B a | B ab | A bc | NS bcd | AB cd | NS d |
|  | Mature Salt | B ab | C a | B ab | B b | NS b | B b | NS b |

**Figure 2.**
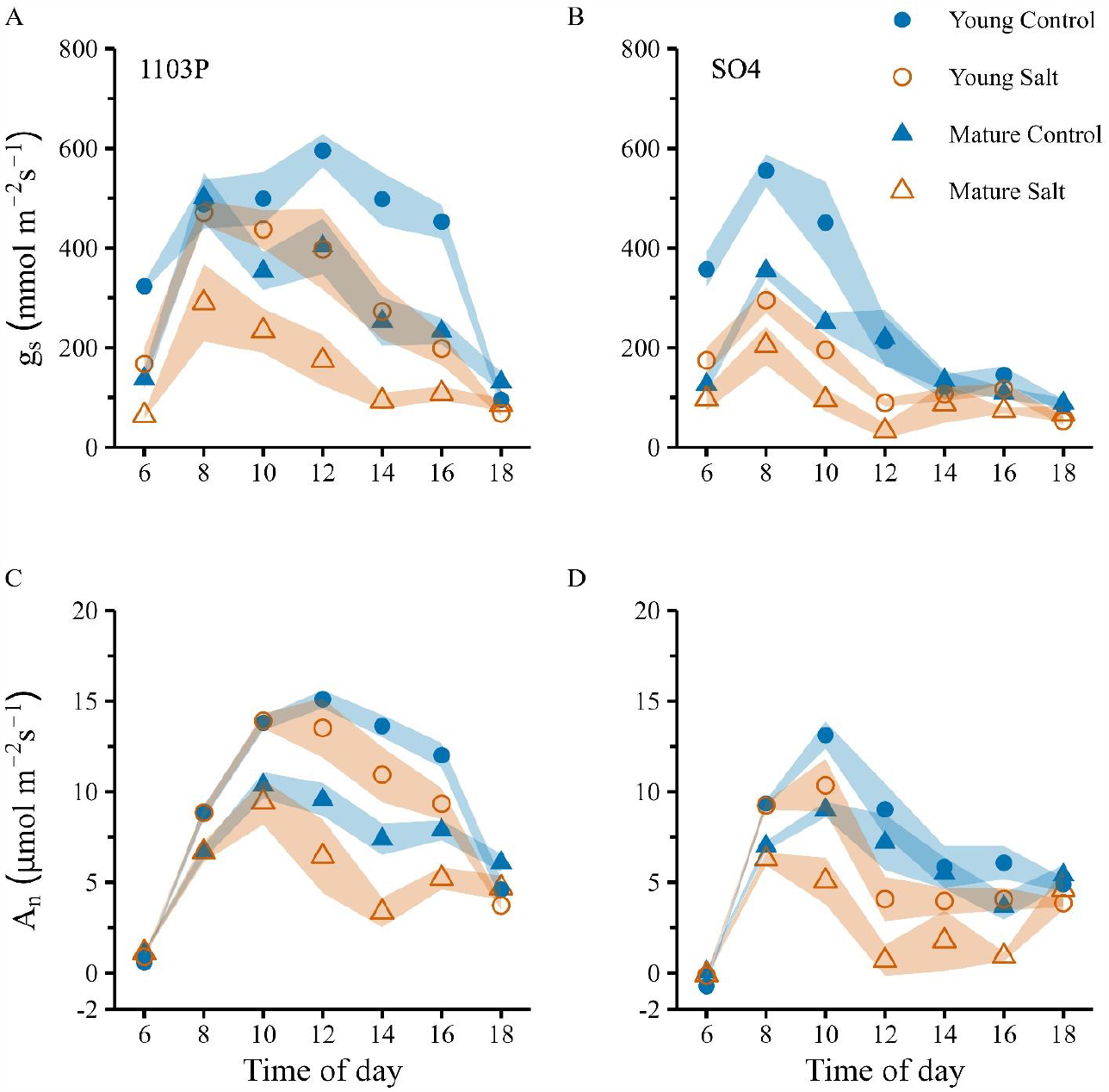
Mean diurnal gas exchange of young (21 days old; circles) and mature (63 days old; triangles) leaves under control (blue, full symbols) and salt (orange, empty symbols) treatments. (a) 1103P’s stomatal conductance (g_s_). (b) SO4’s g_s_. (c) 1103P’s CO2 assimilation rate (A_n_). (d) SO4’s A_n_. Background color represents standard error. Statistical analysis results are presented in table 1 and table 2.

1103P’s young leaves had a diurnal peak of A_n_ at 12 pm under the control treatment, while under the salt treatment, the peak was at 10 am (Fig. 2c, table 2). The daily peak in 1103P’s A_n_ for the mature leaves was at 10 am under both the control and salt treatments. SO4’s young leaves had earlier daily A_n_ peaks compared to 1103P, at 10 am under both the control and salt treatments (Fig. 2d, table 2). SO4’s mature leaves peak was even earlier under the salt treatment, at 8 am. SO4’s leaves lowered their A_n_ during the midday (12 pm), compared to the morning peaks. 1103P’s young leaves had higher A_n_ than SO4 under the control and salt treatments during the midday (student’s t-test, P < 0.05; data not shown), while no significant differences were observed in the mature leaves.

**Table 2.** Two-way ANOVA analysis for the diurnal measurements of A_n_. Results are presented in figure 2c, d. Different uppercase letters represent significant differences between leaf ages and treatments within each time point. Different lowercase letters represent significant differences between time points within each leaf age-treatment combination. Tukey HSD, P < 0.05. n = 6.

| Variety | Treatment/Hour | 06:00 | 08:00 | 10:00 | 12:00 | 14:00 | 16:00 | 18:00 |
| --- | --- | --- | --- | --- | --- | --- | --- | --- |
| 1103P | Young Control | B e | A c | A ab | A a | A ab | A b | AB d |
|  | Young Salt | A c | A b | A a | A a | AB ab | AB b | B c |
|  | Mature Control | A d | B c | B a | AB ab | B bc | BC abc | A c |
|  | Mature Salt | A c | B ab | B a | B ab | C bc | C abc | AB bc |
| SO<sub>4</sub> | Young Control | B d | A b | A a | A b | NS bc | A bc | NS c |
|  | Young Salt | A c | A a | A a | AB b | NS b | A b | NS b |
|  | Mature Control | A d | B abc | AB a | A ab | NS bc | AB c | NS bc |
|  | Mature Salt | A c | B a | B ab | B bc | NS bc | B bc | NS ab |

### 3.3 Sodium and chloride contents in leaves of different ages

Leaf Cl^-^ contents increased with leaf age under the salt treatment for 1103P and SO4 (Fig. 3a, b). 1103P had higher Cl^-^ contents under the salt treatment in all leaf ages compared to the control. Under the salt treatment, Cl^-^ increased in 1103P with leaf age and was significantly higher in 49 and 63 days old leaves compared to 7 days old (Fig. 3a). SO4 had higher Cl^-^ contents under the salt treatment in all leaf ages, except for 7 days old leaves, compared to the control. Under the salt treatment, Cl^-^ significantly increased in SO4 with leaf age from 7 to 49 days old leaves (Fig. 3b). 1103P had higher Cl^-^ in 21 days old leaves compared to SO4 (Student’s t-test, P < 0.05). Leaf Na^+^ contents in 1103P and SO4 were lower than the Cl^-^ contents and were significantly higher under the salt treatment only in 63 days old leaves compared to the control (Fig. 3c, d).

**Figure 3.**
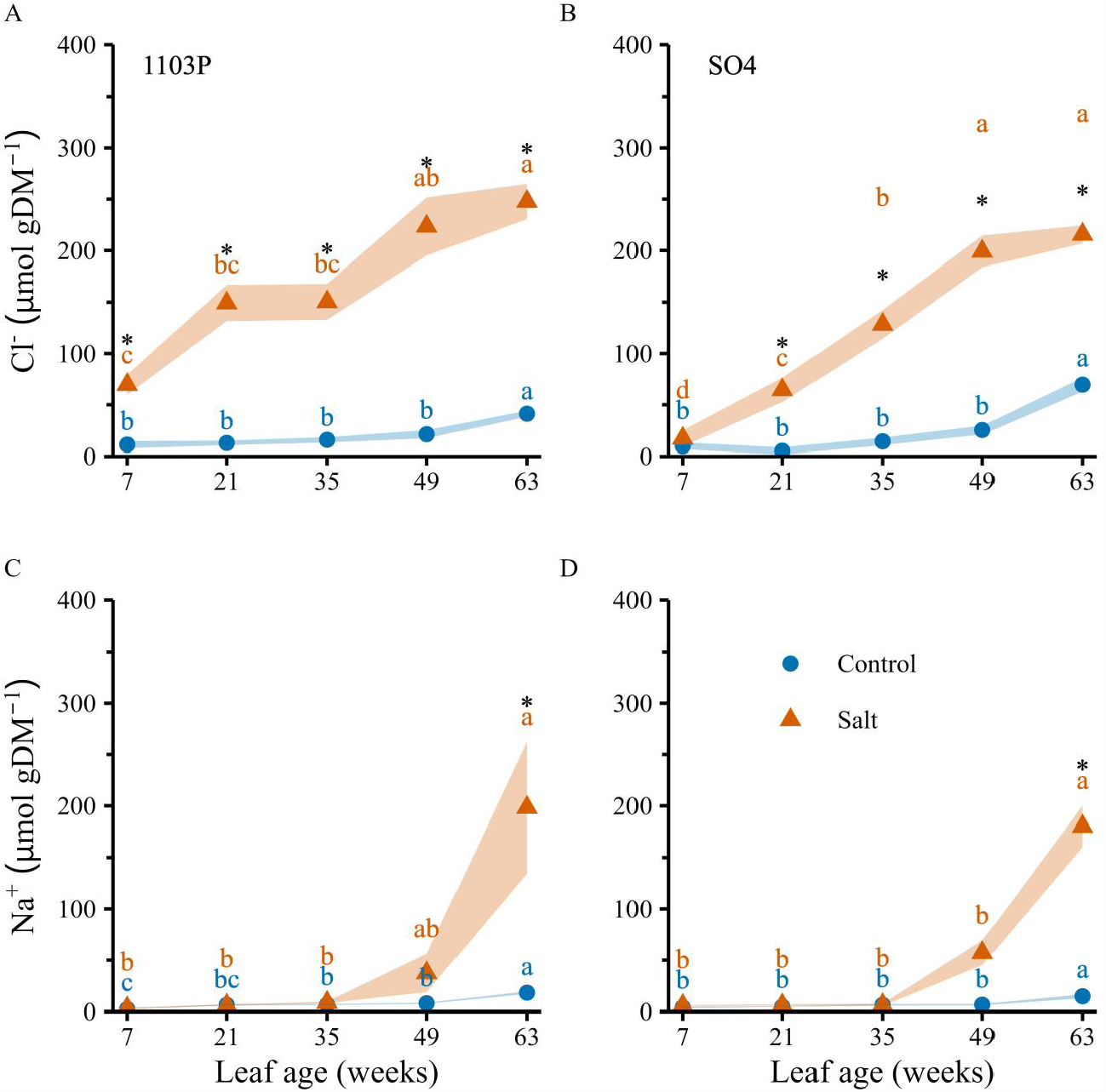
Mean chloride (Cl^-^) and sodium (Na^+^) contents in different leaves ages of 1103P (left) and SO4 (right) under control (blue circles) and salt (orange triangles) treatments. (a) 1103P’s Cl^-^ contents. (b) SO4’s Cl^-^ contents. (c) 1103P’s Na^+^ contents. (d) SO4’s Na^+^ contents. Background color represents standard error. Different letters represent significant differences between leaf ages within each treatment, one-way ANOVA, P < 0.05. Asterisks represent significant differences between treatments within each leaf age, student’s t-test, P < 0.05. n = 6.

### 3.4 Osmolality and carbon to nitrogen distribution in young and mature leaves

Under the salt treatment, the morning osmolality was higher in 1103P’s young and mature leaves and in SO4’s mature leaves, compared to the control (Fig. 4a). SO4 had higher morning osmolality compared to the young leaves under the control and salt treatments while 1103P had higher osmolality only under the salt treatment. SO4 also had higher morning osmolality in its mature leaves under the salt treatment compared to 1103P (student’s t-test, P < 0.05). The midday osmolality levels were generally higher than those in the morning (Fig. 4b). Midday osmolality was higher than morning osmolality at a range of 13.6% (SO4-young-control) to 44.5% (1103P-mature-control). 1103P had higher midday osmolality in its mature leaves under the salt treatment compared to the control. SO4 had higher midday osmolality in its mature leaves compared to the young leaves under the control and salt treatments. SO4 also had higher mature leaves’ midday osmolality than 1103P under the control treatment (student’s t-test, P < 0.05). There were no significant differences in young leaves’ morning and midday osmolality between the varieties under the control and salt treatments.

**Figure 4.**
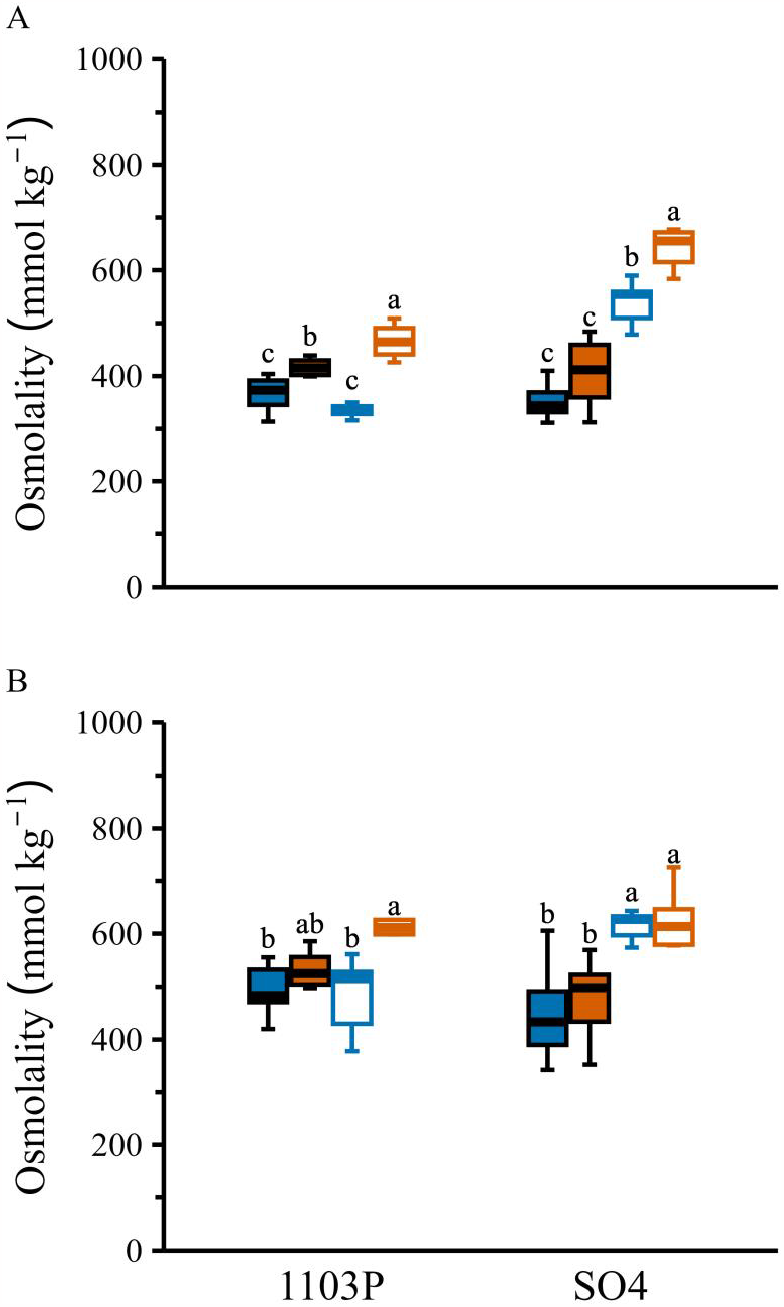
Osmolality of young (21 days; full boxes) and mature (63 days; empty boxes) leaves under control (blue) and salt (orange) treatments. (a) morning (6 am) osmolality. (b) midday (12 pm) osmolality. Different letters represent significant differences between leaf ages and treatments within each variety, two-way ANOVA followed by Tukey HSD, P < 0.05.

The carbon to nitrogen (C:N) ratio was higher in 1103P’s mature leaves compared to its young leaves under the control and salt treatments, while no significant differences were observed in SO4 (Fig. 5a). 1103P had lower C content in its mature leaves than its young leaves (Fig. 5b) under the salt treatments and lower N content under the control and salt treatments (Fig. 5c). SO4 had higher C content in its mature leaves than its young leaves under the salt treatments (Fig. 5b), but had no significant differences in N between leaf ages and treatments (Fig. 5c). There were no significant differences in shoot mass between 1103P and SO4 under the control and salt treatment (Fig. 6a). Both varieties had lower shoot mass under the salt treatment compared to control. 1103P had a higher stem diameter compared to SO4 under both the control and salt treatments, and both varieties had lower stem diameters under the salt treatment compared to the control (Fig. 6b).

**Figure 5.**
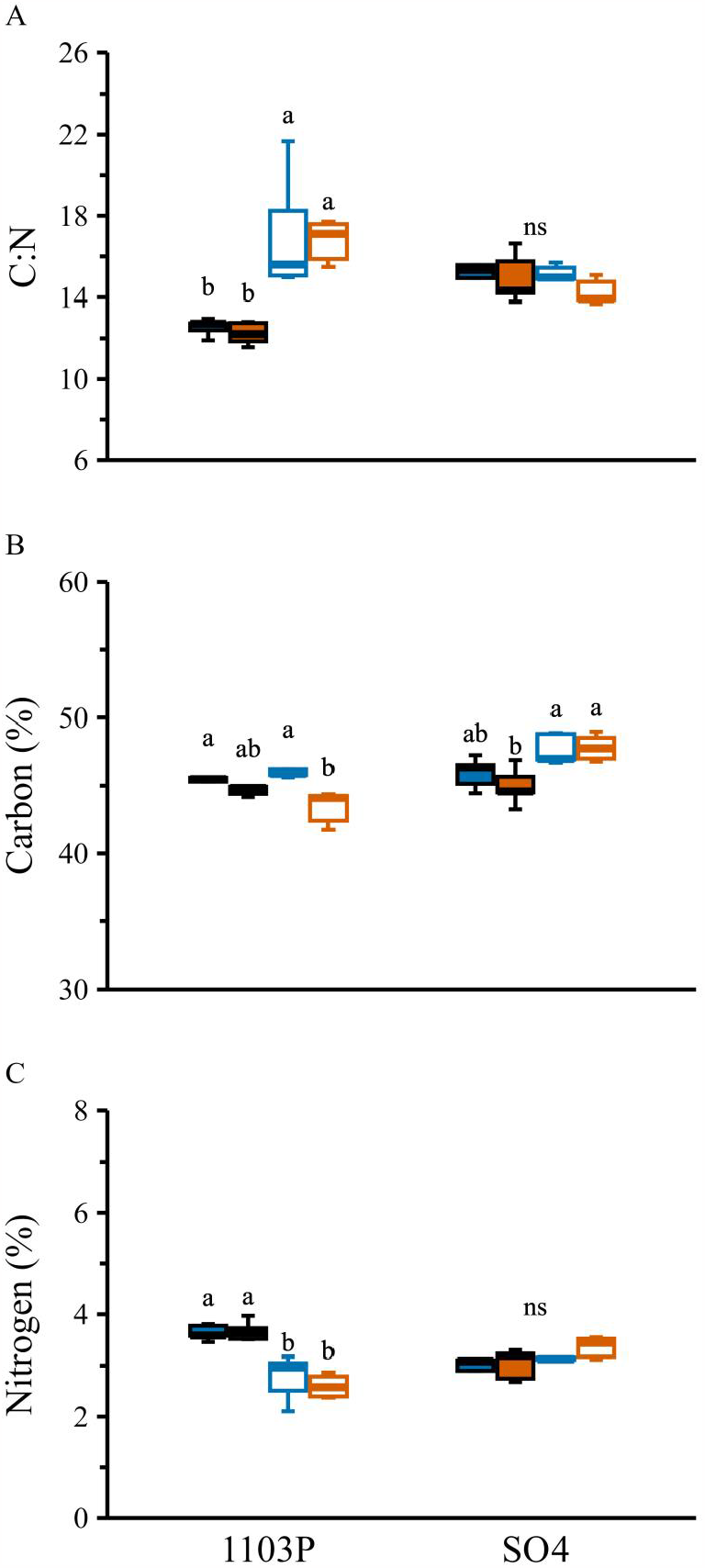
Carbon (C) and nitrogen (N) contents of young (21 days; full boxes) and mature (63 days; empty boxes) leaves under control (blue) and salt (orange) treatments. (a) carbon to nitrogen ratio (C:N). (b) carbon contents. (c) nitrogen contents. Values are a percent of dry weight. Different letters represent significant differences between leaf ages and treatments within each variety (ns = non-significant), two-way ANOVA followed by Tukey HSD, P < 0.05.

**Figure 6.**
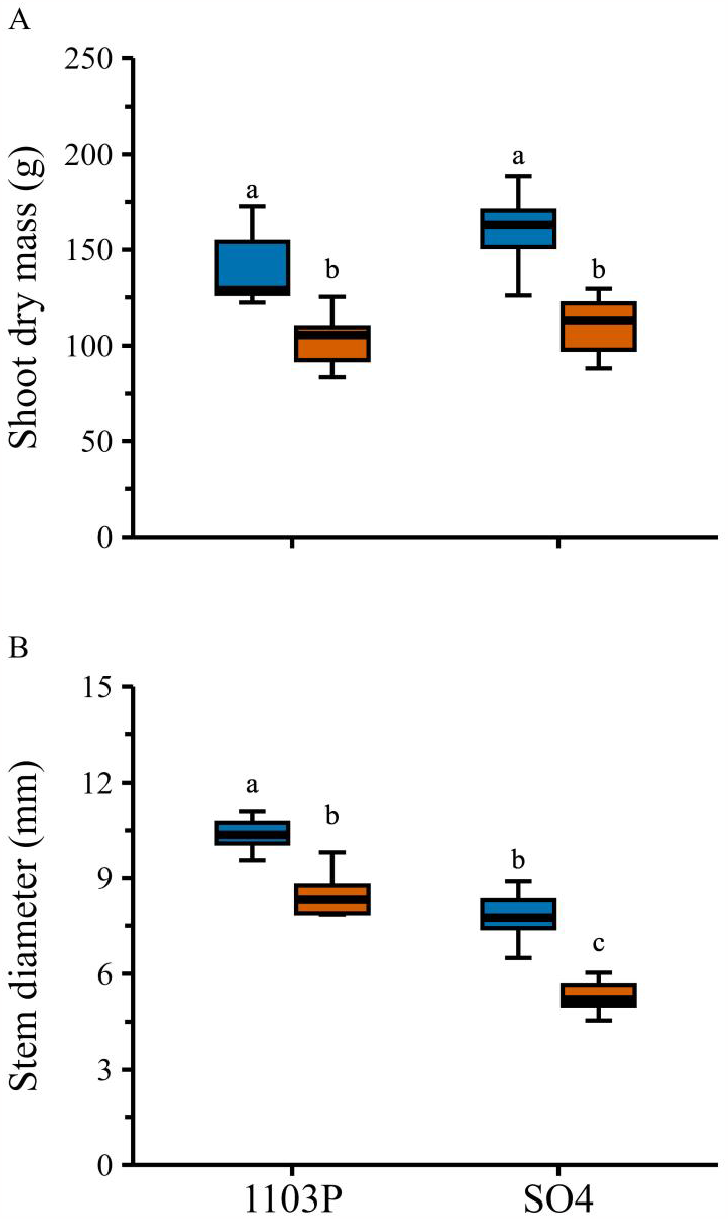
Shoot dry mass and stem diameter at the end of the experiment under control (blue) and salt (orange) treatments. (a) Shoot dry mass. (b) stem diameter. Different letters represent significant differences between varieties and treatments, two-way ANOVA followed by Tukey HSD, P < 0.05. n = 9.

## 4. Discussion

Leaf age highly affected A_n,_ as it reached a maximum when leaves were 28 days old and declined as leaves matured (Fig. 1c, d). However, g_s_ was minimally affected by leaf age. This leaf age-associated decline in A_n_ was not due to a decrease in g_s_ but was partially a result of declining PSII efficiency (Fig. 1e, f). In contrast, salt stress lowered g_s_ but had a low impact on A_n_ (Fig. 1a, b). We also found that leaf age had a differential effect on Na^+^ and Cl^-^ exclusion capacities (Fig. 3). Na^+^ did not accumulate in the leaves until they were mature (63 days old), while Cl^-^ started to accumulate at very young leaves (7 days old) and continued to accumulate as the leaves matured. Additionally, mature leaves had higher osmolality due to increased salt accumulation rather than osmotic salt stress adjustments (Fig. 4). These findings partially support our hypothesis that salt concentration would increase with leaf age and be associated with decreased leaf function, as salt concentration increased with leaf age but did not directly affect leaf performance.

### 4.1 Leaf age-associated decline in photosynthesis is initially affected by a reduction in PSII

Photosynthesis was correlated with leaf age (P < 0.01) in both varieties. The photosynthetic rate showed similar trends, with an increase as the leaves matured to a maximum at leaf age of 28-35 days following a constant decline (Fig. 1c, d). This trend is not similar for all crops, as maize, a C4 annual cereal, had maximum photosynthesis before the leaf was fully expanded, followed by a linear decrease (Dwyer and Stewart, 1986). The age-associated decline in A_n_ cannot be entirely attributed to a reduction in g_s_, as it was less affected by the leaf age and did not follow a similar pattern as A_n_ (Fig. 1). Therefore, other factors play a role in the age-associated A_n_ decline. An increase in chlorophyll content might explain the increase in A_n_ between the young (14 days old) and young fully expanded (28 days old), as chlorophyll was found to be low in young grapevine leaves (Bertamini and Nedunchezhian, 2003). It was suggested that age-associated decline in photosynthesis results from lower chlorophyll content and lower Rubisco activity in 90 days or older leaves (Bertamini and Nedunchezhian, 2002). We found significant age-associated photosynthesis declines in leaves at 42-49 days, suggesting that more factors are involved. One such factor is PSII efficiency which is the first step in the light-dependent phase of photosynthesis. A_n_ followed a similar reduction trend of age-associated declines in PhiPS2 (Fig. 1; an average of -16.7% in A_n_ and -13.85% in PhiPS2 between 28 and 42 days old leaves). Thus, PSII is the first photosynthesis compound to decline as the leaves mature. Therefore, it is likely that the age-associated decline in photosynthesis starts with a reduction in the light-dependent phase (PSII) followed by a reduction in the light-independent phase (rate of carboxylation).

### 4.2 Fast closure of the stomata under salt stress leads to improved WUE but with a tradeoff in plant growth

While the salt stress had a low to no effect on A_n_, the g_s_ was lowered under salt stress for both varieties when the leaves were younger than 56 days (Fig. 1). Similar results were found in grapevines, in response to drought, as g_s_ was reduced by 46% while the A_n_ was reduced by only 16% (Poni et al., 2007). Photosynthesis is considered less sensitive to turgor (Taiz et al., 2015), thus explaining how the salinity-induced osmotic stress reduced the plants’ g_s_ but not A_n_. The few cases A_n_ was found to reduce in response to salt stress (SO4’s 28, 42 and 49 days old leaves) can be attributed to lowered g_s_ as the Ci was also significantly reduced (23%, 6%, and 12% lower than the control; student’s t-test, P < 0.05; data not shown). As it turns out, g_s_ responses faster than A_n_ to environmental changes allowing to maintain photosynthesis under unfavorable conditions. The salt-induced reduction in stomatal conductance was mainly osmotic, not ionic, as it was not associated with the accumulation of salts in the leaves. The reduction in g_s_ without a following reduction in A_n_ in response to salt stress resulted in an improved iWUE for both varieties, except for SO4’s 28 days old leaves. SO4, which had stronger reductions in g_s_ under salt stress, also had higher iWUE than 1103P. This improvement of WUE under stress was previously reported in grapevines under drought stress (Chaves et al., 2007). High WUE is desirable for crop plants under limiting and beneficial conditions, but only if it comes without reducing productivity (Blum, 2005). Here we show that it is not the case as the shoot mass and stem diameter were reduced under saline conditions (Fig. 6). Different approaches were suggested to improve WUE without a tradeoff in productivity (Flexas et al., 2013; Yang et al., 2019). Therefore, in the face of present and future climate changes in traditional grapevine growing areas (Gambetta and Kurtural, 2021; Morales-Castilla et al., 2020), increased WUE without tradeoff in yield under stress conditions should be considered in future viticulture breeding programs.

### 4.3 Young and mature leaves’ responses to diurnal changes are variety depended and affect the whole-day iWUE

1103P and SO4 had different diurnal water balance trends under beneficial conditions (i.e., young leaves under the control treatment). SO4 was shown to also have a more plastic response to salt stress (Lupo et al., 2022). SO4’s diurnal trend of high g_s_ in the morning followed by a decline in the midday enables it to be more efficient in its water use, having 28 % higher daily iWUE than 1103P (data not shown). This trend of morning g_s_ peak was also found in tomatoes under good irrigation and was correlated with high yield (Gosa et al., 2022). The differences in midday g_s_ between 1103P and SO4 were not a result of differential osmolytes accumulation, as there were no significant differences in midday osmolality between them (two-way ANOVA, P < 0.05; Fig. 4b). When considering 6 am, as a base point in which VPD is relatively low and not limiting, the age-associated decline in g_s_ was 57% and 65% for 1103P and SO4, respectively. Thus, each day in the leaf age lowered the g_s_ potential by 1.5% on average. Age-associated decline in A_n_ did not appear at 6 am, probably because not all the PSII and PSI reaction centers are activated at this time. However, at 8 am, SO4 had similarly age-associated lower g_s_ and A_n,_ but 1103P did not, showing that more factors affect photosynthesis and that these effects are variety depended.

### 4.4 Grapevines improved sodium exclusion from the leaves is diminished when leaves mature

Unlike most crop plants, Grapevines are more sensitive to Cl^-^ than Na^+^ since they are good sodium excluders (Munns and Tester, 2008). Our data show that while Cl^-^ increased with leaf age, Na^+^ started to increase only when the leaves matured (Fig. 3). Grapevines’ reasonable leaf Na^+^ exclusion is attributed to mechanisms that transfer and store Na^+^ in the woody stems (Walker et al., 2014). How SO4 better excluded Cl^-^ from its young leaves is yet to be uncovered and might arise from having high root’s Cl^-^ transporters activity (Henderson et al., 2014) as SO4 was also found to better exclude Cl^-^ from its roots (Lupo et al., 2022). Our results suggest that the grapevines’ Na^+^ exclusion mechanisms reduce their function when leaves are over 49 days old. The accumulation of Cl^-^ in the leaves did not adversely affect photosynthesis as the salt-treated plants followed a similar trend of age-associated decline in A_n_ as the control plants. The salt stress-induced reduction in g_s_ is likely due to osmotic rather than ionic stress since the increase in leaf Cl^-^ and Na^+^ did not correlate with g_s_. The higher osmolality of the mature leaves did not reflect adaptive aspects in the diurnal g_s._ Therefore, it is more likely attributed to salt accumulation rather than organic osmolytes (Munns et al., 2020). 1103P’s high C:N ratio due to lowering nitrogen content in its mature leaves, under the control and salt treatments, is most likely due to senescence mechanisms that enhanced under salt stress and can also be seen in its salt-induced leaf chlorosis symptoms (S Fig. 1b; Shapira et al., 2009).

### 4.5 Measurements of only the young, fully expanded leaves do not reveal the whole plant’s status

Examining only the youngest, fully expanded leaves for leaf gas exchange and ion content needs to be re-evaluated. The midday gas exchange measurement shows that for SO4, measuring the young leaves might be a good indicator of the whole plant’s status. However, for 1103P, this is not the case since the old leaves had 32% lower g_s_ and 37% lower A_n_ than the young leaves. Moreover, at 49 days old, leaves still perform relatively high photosynthesis (∼60% from maximum A_n_). The same goes for leaf ion content. As salt exclusion from the shoot is a salt-tolerant indicator, examining only young leaves (21 days old), we can say that SO4 is a better excluder than 1103P. However, examining leaves just 14 days older shows no differences between the varieties. The decision of which leaf to sample also has agricultural importance since a major proportion of grapevine foliage consists of leaves older than 21 days for most of the growing season, and leaves can be more than triple this age when harvest starts. Therefore, we suggest that sampling leaves of different ages will be more insightful when looking for a salt-tolerant plant or salinity-caused damage to a plant.

## 5. Conclusions

Leaf age had a significant impact on photosynthesis but only a minimal effect on stomatal conductance. Meanwhile, salt stress had a low impact on photosynthesis but significantly reduced stomatal conductance. The decline in photosynthesis associated with leaf age began with a reduction in PSII efficiency and was not a result of lower stomatal conductance or ionic stress as it was not associated with the accumulation of salts in the leaves. Our findings suggest that while g_s_ was less affected by leaf age, the stomata closed in response to the salt stress, preventing the adverse effects of the osmotic stress by not lowering photosynthesis. Additionally, our study revealed that grapevines’ ability to exclude Na^+^ decreased with leaf maturity, reaching a similar level to Cl^-^ when leaves were sixty-three days old. When assessing plants’ salt tolerance, it is essential to consider leaf age and variety, as gas exchange and salt accumulation depend on both. Therefore, it is essential to sample both mature and young leaves to comprehensively understand the plant’s response to stress and avoid drawing incorrect conclusions.

## Abbreviations

1103P: 1103 Paulsen
SO4: Selection Oppenheim 4

## Acknowledgments

We thank Mr. Danny Wais from the “Hishtil” nursery for their kind donation of rootstocks cuttings. This work was supported by the Israeli ministry of agriculture and rural development as part of ERA-NET 5317. This research was part of the EnViRoS project (Opportunities for an Environmental friendly Viticulture: optimization of water management and introduction of new Rootstock and Scion genotypes).

