## Supplementary material for "Importance of Leaf Age in Grapevines Under Salt Stress": S Fig. 1

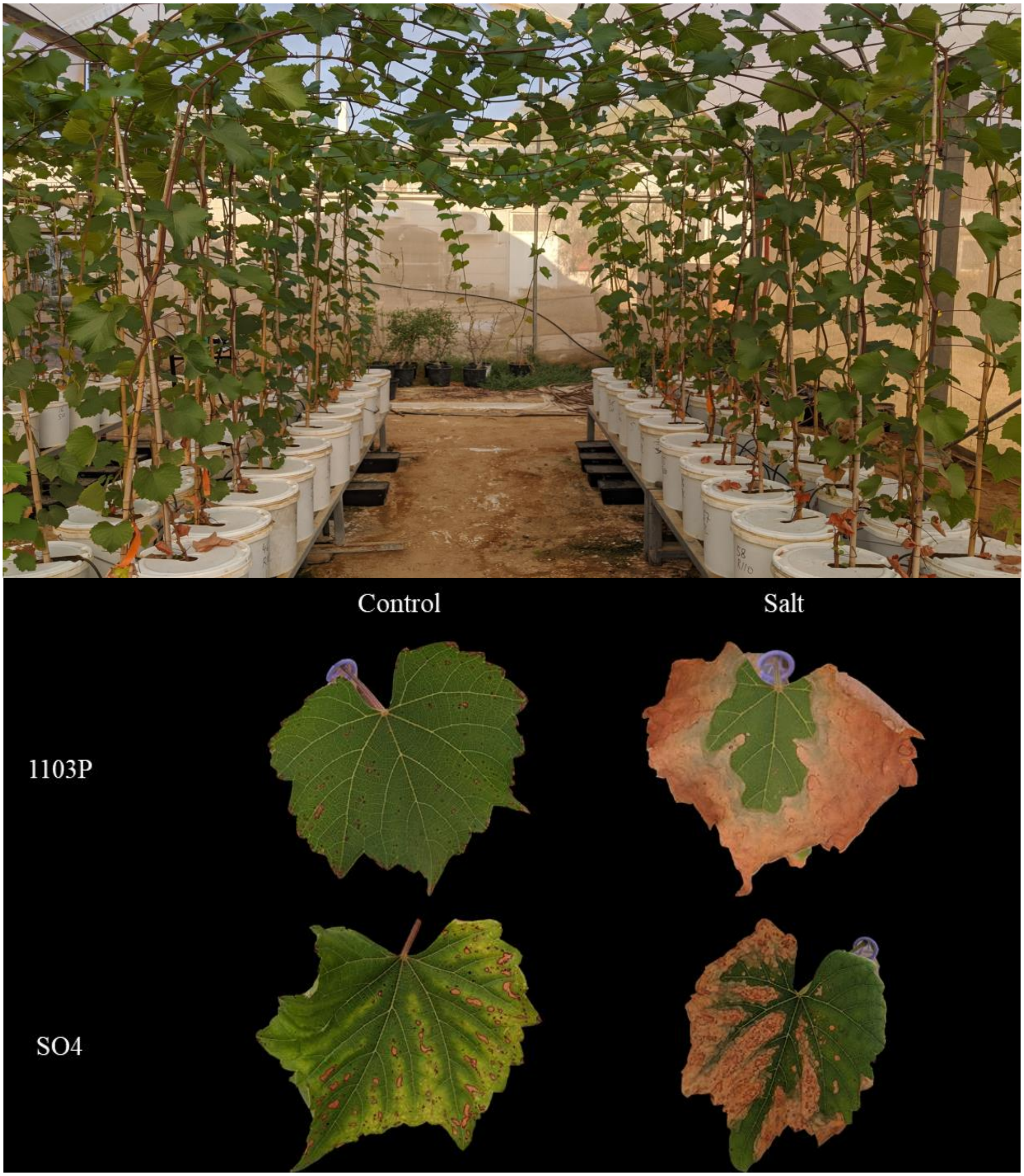


**Supplementary figure 1.** (a) experimental setup at the end of the experiment. (b) 77 days old leaves under the control and salt treatments.
